## Supplementary for "Reductions in prefrontal activation predict off-topic utterances during speech production"

### Supplementary Methods

List of prompts used to elicit speech:

Which is your favourite season and why?

What do you like or dislike about Christmas?

Why do people come to Scotland on holiday?

What sort of things usually happen at a wedding?

Describe a typical visit to a restaurant

What sort of things do you have to do to look after a dog?

What do people usually do when getting ready for work in the morning?

Describe the steps you would take if going somewhere by train

Describe how you would make a cup of tea or coffee

How would you prepare to go on holiday?

What would it have been like to live in the Middle Ages?

What would it be like to live in Antarctica?

What do the police do when a crime has been committed?

What happens when a storm is forecast in the UK?

What are the advantages and disadvantages of going to university?

What sort of things does the Queen do on a typical day?

What happens during a general election in the UK?

Why are some people concerned about climate change?

Do you think it's a good idea to send people to live on Mars?

Do you think the internet has improved people's lives?

Further methods of LSA space and averaging method: The LSA vector space was generated using the British National Corpus [1]. Each document in the BNC was divided into contexts of 1000 words in length. A term-by-context matrix was generated, log-entropy weighting applied and singular value decomposition used to reduce the dimensionality of the matrix to 300 dimensions. These steps were performed using the Text to Matrix Generator toolbox in Matlab [2] and resulted in the creation of latent semantic vectors for 53,758 words. To generate semantic representations for speech windows or whole responses, the vectors for individual words were combined in the following way:

1. A list of words appearing in the speech window was created. Common function words that carry little semantic information were removed from the list.
2. Vectors for each word in the list were retrieved and normalised so that each had a magnitude of one.
3. The vector for each word was weighted according to (a) the log of its frequency in the speech passage being analysed (assigning greater weight to words occurring more frequently in the window) and (b) its entropy value in the BNC (assigning greater weight to words whose presence is more informative about the semantic content of the window).
4. The weighted vectors were averaged to give a single vector representing the semantic content of the words in the window.

These procedures were similar to those used by other researchers [3,4].

Measurement of other characteristics of speech: Transcripts were submitted to the Stanford Log-linear Part-of-Speech Tagger v3.8 for automated part-of-speech tagging [5]. In addition to the coherence measures described in the main text, the following measures were computed for each 5s block of speech:

Number of words: the total number of words produced in the 5s speech block.

Proportion closed-class words: The proportion of words in each block whose part-of-speech was classified as closed-class. Closed-class words included pronouns, numbers, prepositions, conjunctions, determiners, auxiliaries and some adverbs.

Type: token ratio (TTR): This is the ratio of unique lexical items (types) produced to total words (tokens) spoken. A higher value indicates greater lexical diversity. This is the only speech marker that could not be calculated for individual blocks; it was instead calculated for each 50s response as a whole. Because TTR is highly dependent on response length, a moving window approach was adopted to control for the number of words included in the analysis [6]. For each response, the first 50 words were first considered and a TTR calculated over these. The window was then moved forward one word and a new TTR calculated, and this process was repeated until the end of the response was reached. A mean TTR was then computed across all windows.

Mean noun frequency: Frequencies in the SUBTLEX-UK database [7] were obtained for all words tagged as nouns and an average calculated for each block.

Mean noun concreteness: Concreteness ratings for nouns were obtained from Brysbaert et al. [8].

Mean noun age of acquisition (AoA): Estimates of AoA for nouns were obtained from the norms of Kuperman et al. [9].

Mean noun semantic diversity (SemD): SemD values for nouns were obtained from Hoffman et al. [1]. SemD is a measure of variability in the contextual usage of words. Words with high SemD values are used in a wide variety of contexts and thus more variable and less well-specified meanings.

Mean noun number of phonemes: The length of all nouns (in phonemes) was also calculated.

Local coherence: Local coherence refers to the degree to which adjoining utterances in speech are meaningfully related to one another. A measure of local coherence was computed using the same methods as Hoffman et al. [10]. Like the main global coherence metric, this was based on LSA analysis of speech produced by each participant. However, rather than comparing each window of speech to the prototypical response to the topic, each 20-word window was compared to the participant's own previous window. Thus, the measure indexed the degree to which current speech production was meaningfully related to utterances produced earlier in the response.

In cases where a particular measure could not be computed for a block (for example, blocks in which no nouns were produced) the mean value for the rest of the response was used.

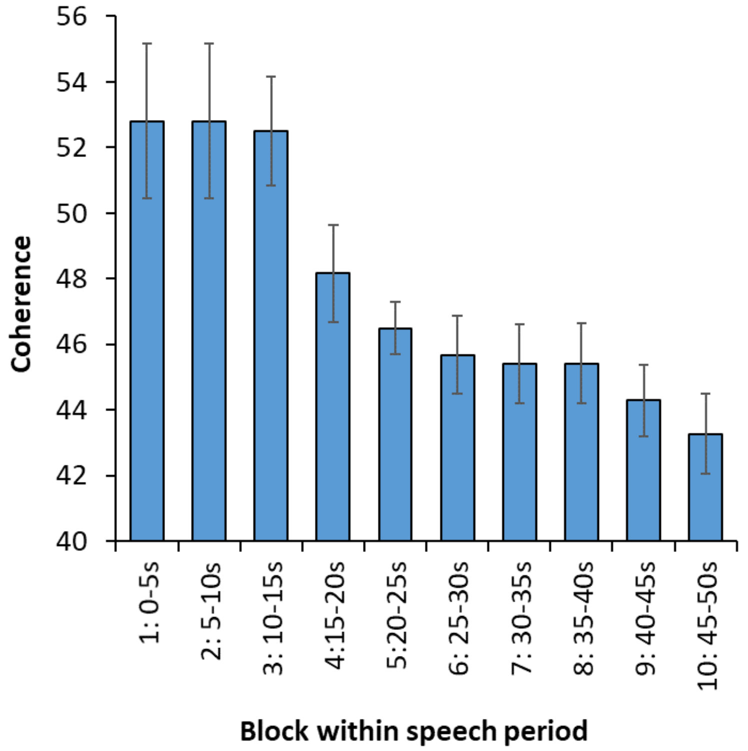


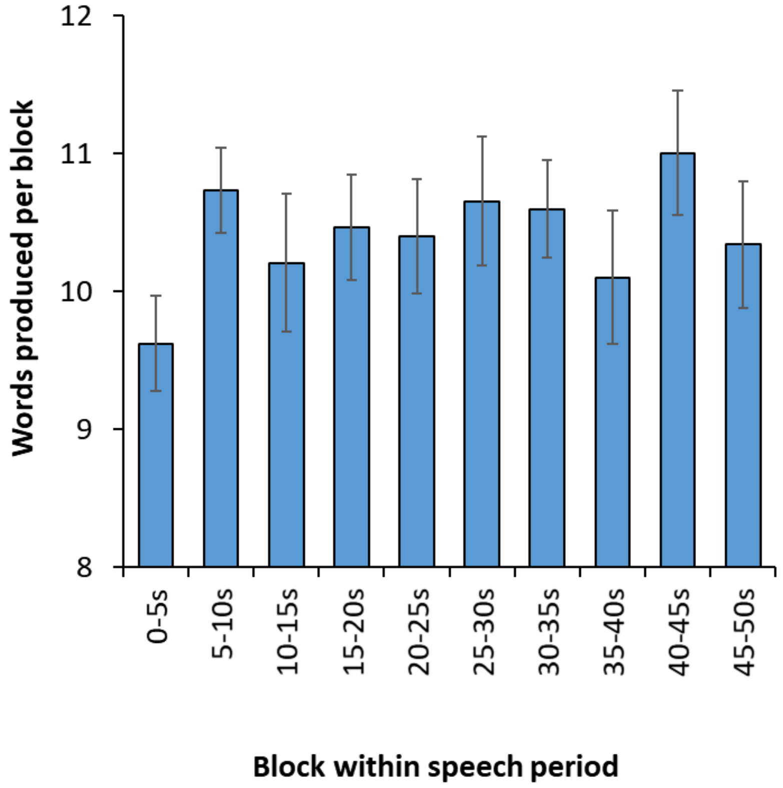


*Supplementary Figure 1: Average coherence of speech and number of words produced in each 5s block of the speech production periods. Bars indicate standard error of the mean.*

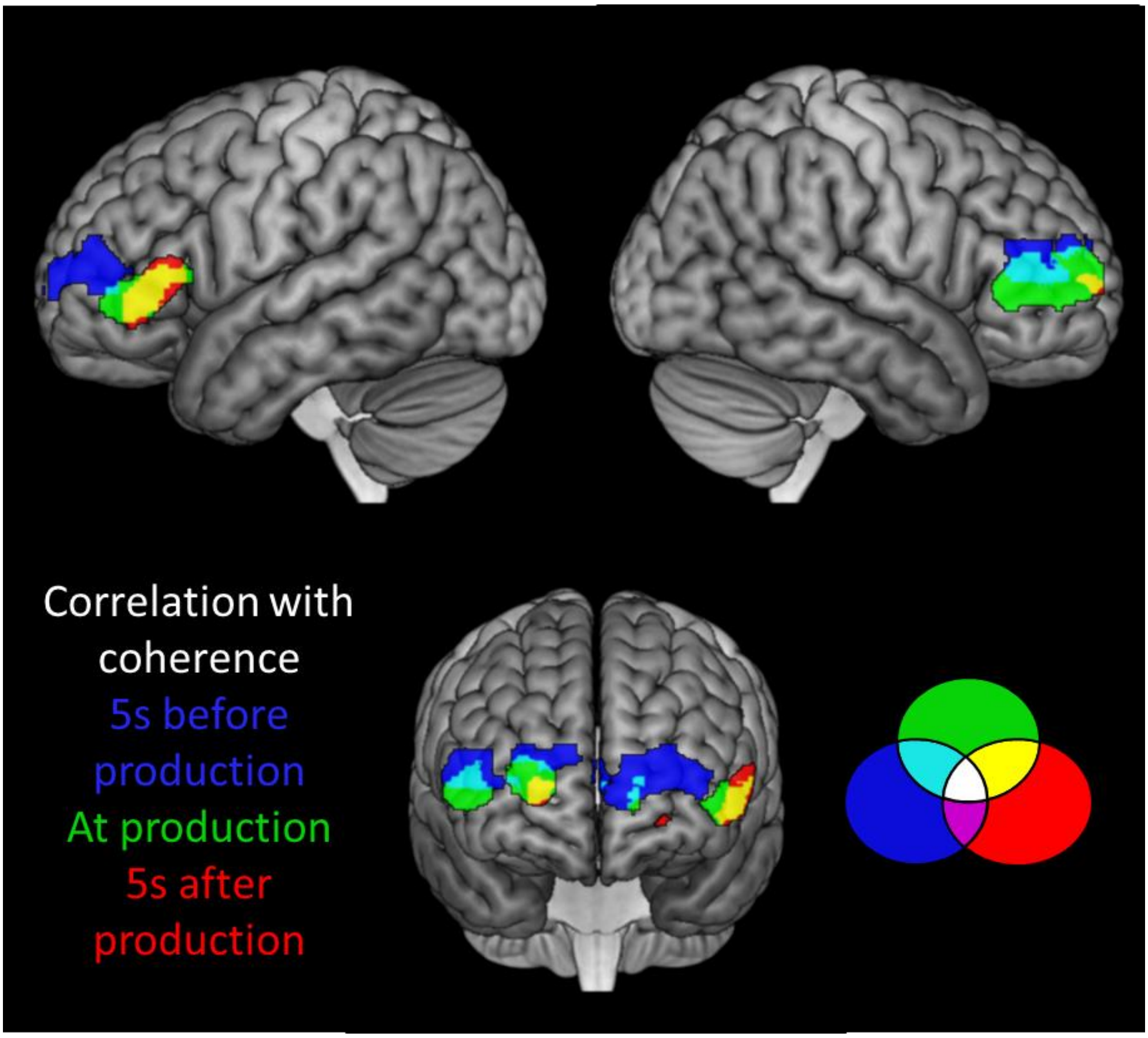


*Supplementary Figure 2: Regions showing greater activation when participants produced more coherent speech at time of production (green; standard model), 5s before the speech was produced (blue; early model) and 5s after the speech was produced (red; late model).*

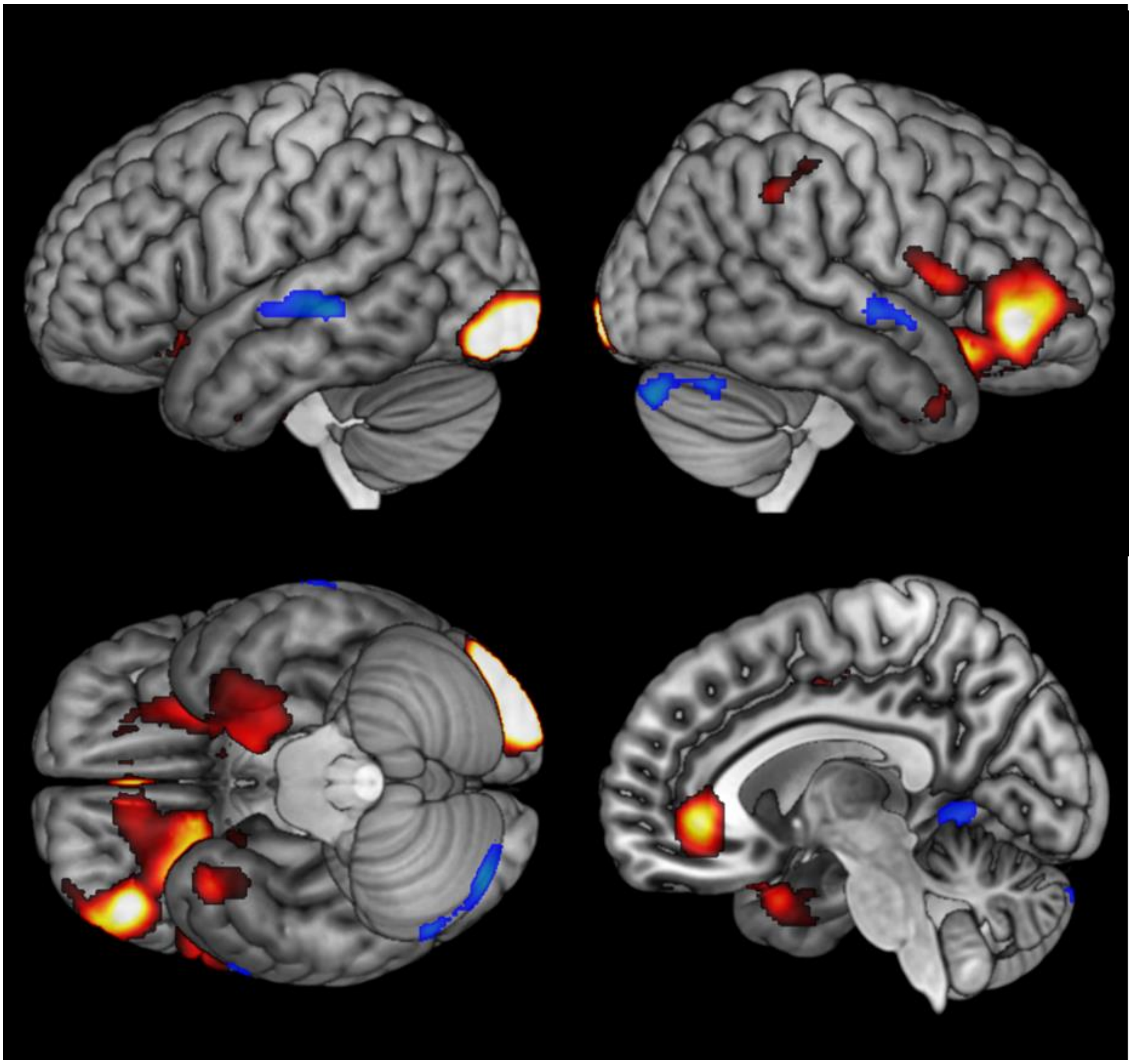

*Supplementary Figure 3: Regions showing increasing (hot colours) or decreasing (cool colours) activation over time within speech production periods.*

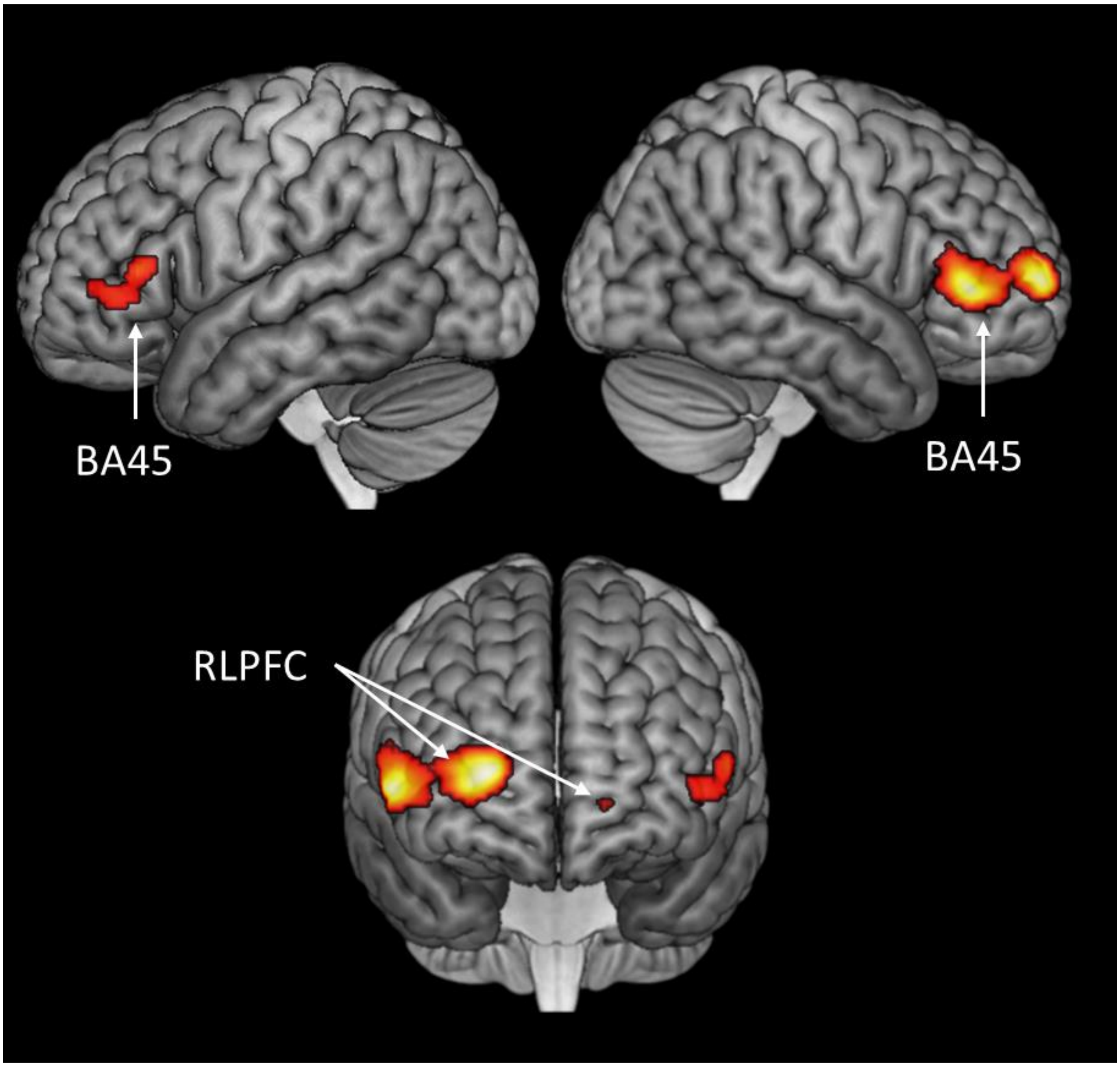


*Supplementary Figure 4: Regions showing a positive correlation with the coherence factor obtained from principal components analysis of speech properties.* *The other PCA factors (corresponding to vocabulary, specificity and verbosity) were entered as covariates in this analysis, thus observed effects cannot be due to covariance with these aspects of speech.*

*Supplementary Table 1: Activation peaks for speech planning and production, relative to baseline*

| Contrast | Location | Extent ($mm^3$) | *t* | MNI co-ords x | y | z |
|---|---|---|---|---|---|---|
| Speech planning | Occipital pole | 95392 | 10.45 | 2 | -94 | -4 |
| | R lingual gyrus | | 6.87 | 16 | -78 | -14 |
| | R cerebellum | | 6.01 | 42 | -66 | -26 |
| | L superior parietal | | 4.45 | -24 | -78 | 54 |
| | L angular gyrus | | 4.27 | -40 | -82 | 38 |
| | R temporal pole | 3544 | 7.41 | 58 | 12 | -24 |
| | R anterior STS | | 4.67 | 68 | -2 | -10 |
| | L anterior MTG | 11256 | 6.76 | -68 | -8 | -14 |
| | L anterior MTG | | 6.53 | -54 | 0 | -26 |
| | L mid MTG | | 3.62 | -56 | -34 | -6 |
| | L pars orbitalis | 23544 | 5.83 | -44 | 34 | -14 |
| | L middle frontal gyrus | | 5.56 | -44 | 6 | 58 |
| | L pars triangularis | | 5.43 | -46 | 24 | 4 |
| | Substantia nigra | 11336 | 5.51 | -8 | -22 | -14 |
| | R globus pallidus | | 5.36 | 16 | 2 | -4 |
| | L globus pallidus | | 5.26 | -16 | 4 | 6 |
| | L SMA | 1264 | 4.03 | -6 | 10 | 58 |
| Speech production | R cerebellum | 22112 | 8.99 | 36 | -78 | -34 |
| | R cerebellum | | 5.18 | 20 | -92 | -30 |
| | R cerebellum | | 4.83 | 16 | -78 | -24 |
| | L lateral occipital | 7192 | 6.60 | -44 | -84 | 32 |
| | L angular gyrus | | 4.20 | -42 | -64 | 18 |
| | L superior frontal gyrus | 7760 | 5.26 | -8 | 32 | 62 |
| | L SMA | | 4.99 | -8 | 18 | 60 |
| | L superior frontal gyrus | | 3.70 | -10 | 50 | 48 |
| | L middle frontal gyrus | 6800 | 5.24 | -40 | 14 | 58 |
| | L middle frontal gyrus | | 5.15 | -36 | 10 | 64 |
| | L pars triangularis | 2376 | 4.99 | -52 | 24 | 6 |
| | L anterior MTG | 7000 | 4.91 | -60 | -8 | -28 |
| | L anterior ITG | | 4.55 | -44 | -8 | -50 |
| | L anterior ITG | | 4.27 | -46 | -8 | -40 |

*Supplementary Table 2: Activation peaks associated with coherence and time during speech production*

| Effect | Location | Extent (mm³) | *t* | MNI co-ords x | y | z |
|---|---|---|---|---|---|---|
| Positive effect of coherence | R frontopolar cortex | 11720 | 7.20 | 22 | 58 | 6 |
| | R pars triangularis | | 5.49 | 42 | 30 | 4 |
| | R anterior MFG | | 5.13 | 48 | 56 | 2 |
| | L anterior cingulate | 11656 | 5.23 | -16 | 40 | 10 |
| | L pars triangularis | | 4.86 | -44 | 40 | -2 |
| | L anterior insula | | 4.60 | -24 | 30 | 6 |
| Negative effect of coherence | No peaks | | | | | |
| Increase over time | L lateral occipital | 10944 | 8.27 | -42 | -90 | -16 |
| | L occipital pole | | 7.41 | -26 | -104 | -6 |
| | R pars orbitalis | 55048 | 5.69 | 44 | 38 | -6 |
| | Ventromedial PFC | | 4.91 | 4 | 38 | -6 |
| | R pars orbitalis | | 4.63 | 30 | 22 | -16 |
| | R temporal pole | | 3.96 | 36 | 10 | -32 |
| | L parahippocampal gyrus | | 3.90 | -14 | -4 | -30 |
| | R middle frontal gyrus | 3680 | 5.09 | 28 | 6 | 32 |
| | R precentral gyrus | | 4.52 | 34 | 0 | 30 |
| | R precentral gyrus | | 3.70 | 26 | -6 | 40 |
| | R angular gyrus | 1040 | 3.60 | 60 | -46 | 40 |
| | R supramarginal gyrus | | 3.22 | 58 | -34 | 48 |
| Decrease over time | L posterior STS | 3264 | 4.04 | -58 | -32 | -2 |
| | R cerebellum | 2136 | 4.03 | 32 | -86 | -30 |
| | R cerebellum | | 3.84 | 50 | -66 | -28 |
| | R superior temporal gyrus | 1040 | 3.63 | 66 | 2 | -6 |
| | R superior temporal gyrus | | 3.52 | 58 | -8 | -2 |
| | R lingual gyrus | 1360 | 3.40 | 14 | -54 | -2 |
| | R lingual gyrus | | 3.35 | 4 | -48 | 0 |
| | Posterior cingulate | | 3.07 | -4 | -46 | 2 |

*Supplementary Table 3: Correlations among properties of speech*

| | 1. | 2. | 3. | 4. | 5. | 6. | 7. | 8. | 9. |
|---|---|---|---|---|---|---|---|---|---|
| 1. Global coherence | -- | | | | | | | | |
| 2. Number of words | -.01† | -- | | | | | | | |
| 3. Type:token ratio | -.10 | -.04 | -- | | | | | | |
| 4. % closed class | -.05 | .24 | -.10 | -- | | | | | |
| 5. Noun frequency | -.10 | .12 | .07 | -.03† | -- | | | | |
| 6. Noun concreteness | .16 | .003† | -.12 | -.02† | -12 | -- | | | |
| 7. Noun age of acquisition | -.07 | -.12 | .04 | .002† | -.49 | -.41 | -- | | |
| 8. Noun semantic diversity | -.18 | .06 | .09 | -.02† | .63 | -.40 | -.15 | -- | |
| 9. Noun phoneme length | -.05 | -.18 | .03† | -.03† | -.42 | -.19 | .50 | -.15 | -- |
| 10. Local coherence | .39 | .04 | -.18 | -.007† | -.10 | .14 | -.06 | -.12 | -.05 |

Speech measures were computed for each 5s block of speech produced in the experiment (N=3000). Due to the large number of observations, most correlation coefficients were significant at $p<0.05$. † indicates coefficients that did *not* reach this threshold.

*Supplementary Table 4: Results of principal components analysis of speech properties*

| | Factor 1 | Factor 2 | Factor 3 | Factor 4 |
|---|---|---|---|---|
| 1. Global coherence | -.03 | -.03 | **.78** | -.12 |
| 2. Number of words | -.17 | .07 | .02 | **.71** |
| 3. Type:token ratio | .01 | .09 | **-.39** | **-.32** |
| 4. % closed class | .06 | -.05 | .02 | **.80** |
| 5. Noun frequency | **-.58** | **.63** | -.03 | -.03 |
| 6. Noun concreteness | **-.56** | **.77** | -.03 | -.06 |
| 7. Noun age of acquisition | **.89** | .08 | .00 | .02 |
| 8. Noun semantic diversity | -.14 | **.87** | .00 | -.02 |
| 9. Noun phoneme length | **.75** | -.06 | -.03 | -.11 |
| 10. Local coherence | .01 | .06 | **.84** | .01 |

Table shows pattern matrix following promax rotation. Loadings with absolute values greater than 0.3 have been highlighted in bold.

*Supplementary Table 5: Peak co-ordinates for analysis of individual differences in coherence*

| Effect | Location | Extent ($mm^3$) | *t* | MNI co-ords x | y | z |
|---|---|---|---|---|---|---|
| Greater activation in more coherent speakers during Planning | R pars triangularis | 5433 | 8.51 | 54 | 34 | 2 |
| | R pars orbitalis | | 4.82 | 40 | 40 | -6 |
| Greater activation in less coherent speakers during Planning | L anterior ITG | 1360 | 4.67 | -52 | -6 | -38 |
| | L anterior ITG | | 3.26 | -52 | -14 | -42 |
| Greater activation in more coherent speakers during Production | L temporal pole | 2192 | 5.93 | -38 | 18 | -46 |
| | L anterior ITG | | 3.39 | -48 | -2 | -50 |
| Greater activation in less coherent speakers during Production | No peaks | | | | | |

*Supplementary Table 6: Demographic information and mean test scores*

| Measure | Score |
| --- | --- |
| Age | 77.5 (8.4) |
| Sex M:F | 6:9 |
| Years of education | 15.0 (2.9) |
| MMSE /30 | 29.5 (0.7) |
| Category fluency (items per category) | 22.3 (3.9) |
| Letter fluency (items per category) | 17.2 (6.7) |
| Trails A errors | 0.1 (0.4) |
| Trails B errors | 1.0 (1.7) |
| Trails A time (s) | 38.5 (13.6) |
| Trails B time (s) | 80.2 (40.4) |
| SCOLP lexical decision (% correct; chance = 50%) | 94.2 (5.2) |
| Synonym judgement (% correct; chance = 25%) | 72.6 (15.2) |

Standard deviations are shown in parentheses. MMSE = Mini-mental state examination [11]. SCOLP = Speed and Capacity of Language Processing test [12]. For the MMSE, Trails and SCOLP tests, participants' scores were compared with published normative data [13-15]. All participants scored above the 10$^{th}$ percentile for their sex, age and education level on all tests.

*Supplementary Table 7: Examples of highly coherent and less coherent responses to the prompt: What do people usually do when getting ready for work in the morning?*

| Participant | Block | Speech | Global Coherence |
|---|---|---|---|
| Participant A | 1 | When people get ready to go to work in the morning, you would | 43 |
| | 2 | normally have set an alarm clock to go off at a time that would … them in enough | 43 |
| | 3 | time to get ready. They would have time for | 63 |
| | 4 | a shower, perhaps a shave, teeth cleaning, | 70 |
| | 5 | hair drying, breakfast. Making the breakfast | 69 |
| | 6 | first of all, of course, hopefully with toast or scrambled eggs or | 50 |
| | 7 | something to start off the day, and a cup of tea or coffee | 50 |
| | 8 | just to settle you down, ready for the day. Many people have to travel | 33 |
| | 9 | to work so they would have to have appropriate time to do so, whether by train | 18 |
| | 10 | coach, car, bicycle or walking | 48 |
| Participant B | 1 | Well it's a matter of getting up, getting out | 77 |
| | 2 | of bed and getting dressed. Washing, shaving, toiletry. | 77 |
| | 3 | Downstairs, breakfast and then | 80 |
| | 4 | make plans for the day, like going to work. | 45 |
| | 5 | Because being aged ninety two and having | 5 |
| | 6 | retired twenty eight years ago, it's a long time since I was personally…. | 10 |
| | 7 | I very much enjoy the life I have | 15 |
| | 8 | without the demands of the call or the urgency | 11 |
| | 9 | of going through to a work situation. I certainly | 19 |
| | 10 | enjoy not having to drive and finish the morning | 53 |
